# Exposure to the organochlorine pesticide cis-chlordane induces ALS-like mitochondrial perturbations in stem cell-derived motor neurons

**DOI:** 10.1101/2025.06.05.658119

**Authors:** Oliver Clackson, Muhammad Reza Hamid, Adithi Wijesekera, Daniel Kulick, Alison L. O’Neil

## Abstract

Amyotrophic Lateral Sclerosis (ALS) is a debilitating and incurable neurodegenerative disease with unsolved etiology. Due to the large proportion of patients lacking direct disease inheritance, understanding the environmental factors that contribute to ALS development is of high priority. Epidemiological studies have implicated pesticides and other environmental exposures as possible contributors to ALS pathogenesis. Recently, our group determined that the organochlorine pesticide cis-chlordane is toxic to human motor neurons in a dose-dependent manner, causing an ALS-like phenotype in culture and animals with a mode of action independent of its known GABA_A_ antagonism. Here, we aimed to characterize downstream motor neuron phenotypes associated with *cis*-chlordane treatment. We performed bulk RNA sequencing, live imaging, immunofluorescent labeling, and real-time metabolic assays on stem cell-derived motor neurons to assess chlordane-associated phenotypes in vitro. We demonstrate that *cis*-chlordane treatment causes a highly altered mitochondrial phenotype in motor neurons, including increased production of reactive oxygen species, decreased OCR and ATP production, and loss of mitochondrial membrane potential. We further implicate *cis*-chlordane as a possible mediator of potent motor neuron damage, with exposure to the pesticide inducing mitochondrial phenotypes akin to those seen in ALS. We suggest that future studies investigating the role of pesticides in ALS development center upon the organochlorine molecules.

## 1. Introduction

Amyotrophic Lateral Sclerosis (ALS) is an incurable neurodegenerative disease marked by significant, rapid-onset motor cortex atrophy resulting in progressive patient paralysis and eventual incapacity to breathe. Despite a large body of research characterizing ALS symptomology, synthesis of complete etiological frameworks has proven challenging. Genetic studies, while highly useful in characterizing a cohort of ALS patients, further demonstrate the disease’s complexity, with just 10-15% of ALS patients harboring disease inheritance (labelled “familial”). In the remaining “sporadic” ALS population, disease pathogenesis is still indefinite, hypothesized as the product of multiple oligogenic risk variants and a nebulous set of environmental and lifestyle factors. [1]

Familial and sporadic ALS patients demonstrate similar clinical phenotypes; thus, it is hypothesized that disease pathology is consistent across both cohorts [2]. However, due to the high proportion of ALS patients lacking disease inheritance, clarifying the external factors that contribute to symptom onset is of high interest and need. Substantial evidence has been collected implicating pesticide exposure as an environmental factor in sporadic ALS development [3-5], a claim bolstered by possible associations of pesticides with other neurodegenerative diseases, such as Parkinson’s disease [6, 7] and Alzheimer’s disease [7, 8]. While few conclusions have been made regarding which pesticides may contribute to ALS phenotypes, as well as their specific etiological impact, a study surveying patient-donated blood specifically found that samples containing the organochlorine pesticide *cis*-chlordane were 9.1 times more likely to have been derived from an ALS patient than a healthy control (odds ratio = 3.87) [9]. Supplementing this finding, one meta-analysis of the longitudinal Agricultural Health Study described a large proportion of ALS patients as having had exposure to organochlorine pesticides [10], while another study surveying cases from the Centers for Disease Control and Prevention National ALS Registry found the presence of organochlorine pesticides in serum, including oxychlordane the metabolite of chlordane, as indicative of ALS risk [11].

The cyclodiene chlordane, formerly sold in emulsions, dust, powders, oil solutions, or sprays as technical chlordane, was first registered in the United States in 1948 and provided a popular mode of pest control in agriculture and construction. In the late 1980s, the compound was banned for outdoor use in the U.S. due to its suggested carcinogenic properties and possible impact on neural development [12]. Despite this, chlordane has persisted in the environment long after its proscription, with its half-life of 10-20 years allowing for stable preservation in air or soil and bioaccumulation in food, fish, and livestock [13, 14]. Analogous to many cyclodienes, chlordane poisons the nervous system of insects through interference with γ-aminobutyric acid (GABA)-mediated signaling, non-competitively antagonizing GABA_A_-linked chloride channels at the distal picrotoxin site [15]. By this mechanism, chlordane and similar cyclodienes induce sudden-onset, convulsive seizures in humans upon absorption through food, skin, or respiration, with acute exposure in large quantities being sufficient to cause death [16]. Recent studies have suggested that chlordane possesses a more complex toxicological profile, demonstrating dysregulation of lipid levels and antioxidant capacity in mice [17] as well as associations with diabetes [18], developmental delays, and metabolic disorders [19]. However, mechanistic analogues for canonical ALS biomarkers are sparse. Ogata et al. demonstrated chlordane-mediated changes in ETC activity and mitochondrial membrane permeability in rat hepatic cells [20], though these findings have scarcely been supplemented. Other organochlorines such as DDT and its metabolite DDE are known to induce mitochondrial dysfunction through ETC uncoupling [21], yet whether chlordane possesses a similar mechanism of toxicity in humans is unknown.

Recently, our research group established that *cis*-chlordane is highly toxic specifically in human induced pluripotent stem cell (iPSC) derived motor neurons, altering action potential dynamics independent of its known mode of GABA_A_ antagonism [22]. To elaborate on these findings and investigate the downstream effects of *cis*-chlordane treatment on motor neuron physiology, we performed live metabolic assays on *cis*-chlordane treated stem cell-derived motor neurons. Herein, we demonstrate that *cis*-chlordane treatment causes a highly altered, ALS-like metabolic phenotype in stemcell derived motor neurons including reduced oxygen consumption and high oxidative stress, suggestive of possible electron transport chain uncoupling and mitochondrial instability.

## 2. Results

### 2.1. RNA sequencing of *cis*-chlordane-treated motor neurons reveals ALS-like transcriptional changes

To assess cellular perturbations in stem cell-derived motor neurons following *cis*-chlordane treatment, we performed RNA-seq on Islet1-GFP motor neurons following pesticide exposure for 16 hours. Following FACS isolation, motor neuron RNA was collected, sequenced, and subjected to differentially expressed gene (DEG) identification. Overall, when comparing treated to untreated motor neurons, 1764 genes were significantly upregulated and 1806 genes were significantly down regulated. (Fig S1) Generally applicable gene set enrichment (GAGE) analysis using the KEGG pathway database was performed, allowing for the identification of broad biological pathways with altered transcriptomic profiles. (Fig S2) Enrichment analysis of *cis*-chlordane treated motor neuron samples indicated strong upregulation of genes involved in ER stress, ER-associated degradation, the unfolded protein response (UPR), and the ER membrane ubiquitin-ligase complex, as well as downregulation of genes involved in axon attraction, repulsion, and outgrowth (Fig S2). The KEGG pathway database was then searched for relevant disease pathways aligning with the observed differential gene expression. Interestingly, this yielded significant enrichment of the canonical “ALS” KEGG disease pathway (p = 0.018), indicating upregulation in ER stress, autophagy, and mitophagy pathways, and downregulation in oxidative phosphorylation and axonal transport pathways (Fig S3), analogous to an ALS phenotype [23].

### 2.2. Cis-chlordane induces ER stress and upregulation of autophagic proteins in motor neurons

To investigate the cause of increased expression of genes related to the UPR, we investigated whether *cis*-chlordane treatment alters the presence of unfolded protein species in WT 1016a motor neurons. Using the PROTEOSTAT Protein Aggregation Assay we observed no significant differences in the presence of unfolded protein between treated and control samples upon normalization to total protein concentration (Fig S4), suggesting that chlordane-associated ER stress is not mediated by increases in unfolded protein species.

A key factor in the amelioration of ER stress is the ubiquitin-proteasome system. Therefore, we next investigated whether *cis*-chlordane impacted proteasome function. A fluorescence-based assay of proteasome activity revealed no significant differences between treated samples and control (Fig S4). Suggesting that the measured UPR was not in response to inhibition of the 26S proteasome.

An alternative to the ubiquitin-proteasome system is autophagy, and our KEGG analysis also demonstrated that, in motor neurons, *cis*-chlordane treatment causes increased expression of autophagic proteins, including the canonical autophagy initiators P62 and LC3. To elaborate on these findings, we performed fluorescence imaging on *cis*-chlordane treated motor neurons stained with LysoTracker, a cell permeable dye that fluoresces upon sequestration in acidic organelles, such as lysosomes. Fluorescence quantification following *cis*-chlordane exposure demonstrated significant increases in average cellular LysoTracker intensity following one- and three-hour *cis*-chlordane treatment periods (Fig 1). This suggests an increase in acidic vesicle content upon *cis*-chlordane treatment, implicating autophagy as a potential moderator of pesticide-induced ER stress.

**Fig 1.**
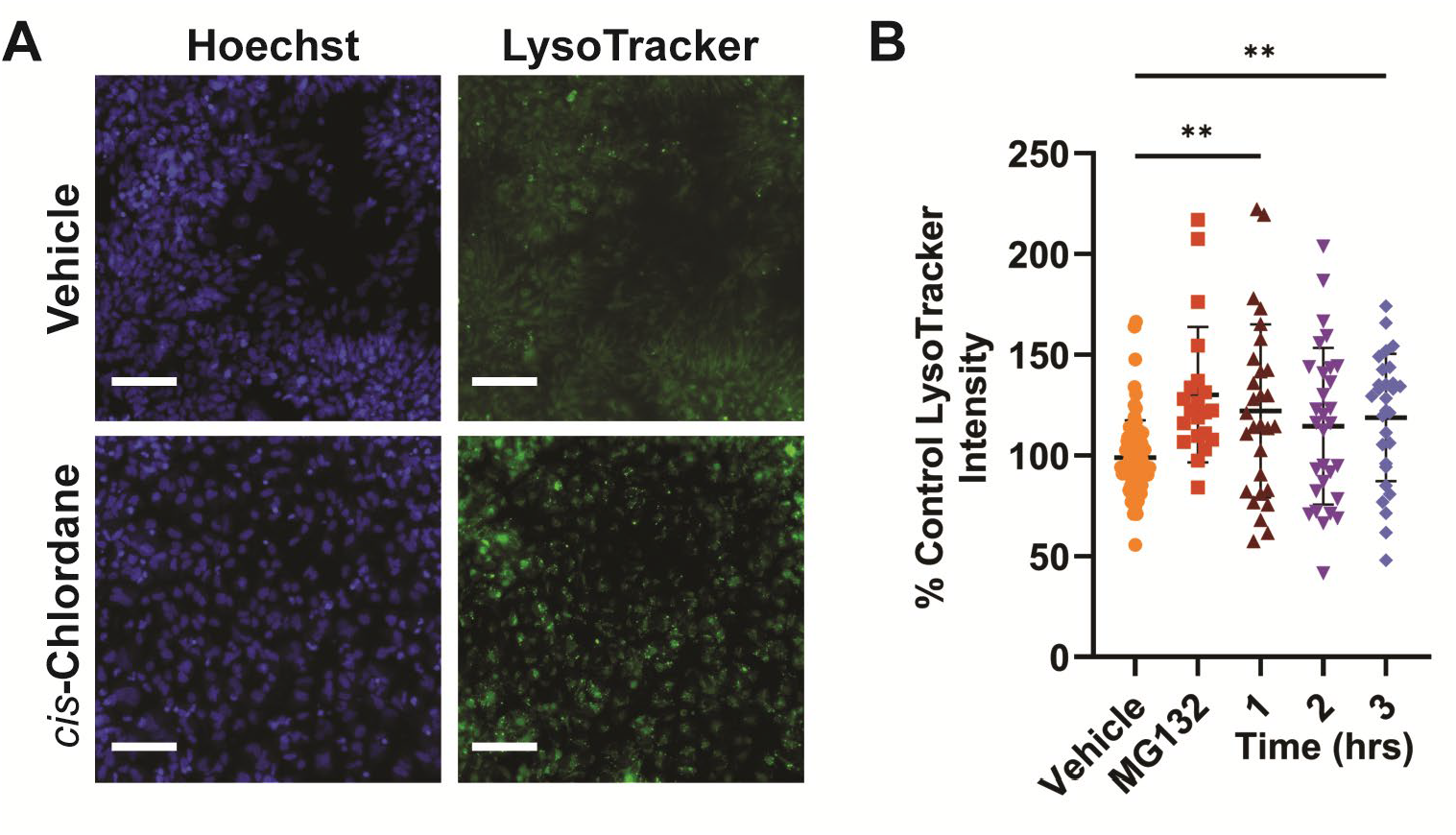
Motor neurons show greater acidic vesicle content following treatment with *cis*-chlordane. Motor neurons were incubated with 10μM *cis*-chlordane, vehicle (DMSO) or the proteasome inhibitor MG132 at 37°C for 1, 2, or 3 hours before staining with LysoTracker dye and subsequent live fluorescent microscopy. (A) Representative live image of motor neurons stained with LysoTracker. Scale bar represents 100µm. (B) Quantification of LysoTracker intensity compared to in-plate negative control. Bars indicate mean with SD. Statistics were performed by Kruskal-Wallis test with multiple comparisons. **p<0.01.

### 2.3. Motor neurons have increased levels of reactive oxygen species in response to *cis*-chlordane treatment

In addition to ER stress, RNA-seq indicated that, in response to *cis*-chlordane exposure, motor neurons decrease expression of genes involved in oxidative phosphorylation and increase mitophagy, suggesting mitochondrial dysfunction. To probe whether motor neurons experience increased oxidative stress following treatment with *cis*-chlordane, we performed fluorescent imaging with CellROX, a cell permeable dye which fluoresces upon oxidation by ROS, as well as MitoSox, a similar dye which accumulates in mitochondria and fluoresces upon oxidation by ROS. After 3 hours of *cis*-chlordane treatment, motor neuron average CellROX intensity was significantly increased, suggesting ROS accumulation in the cytosol in response to pesticide exposure. (Fig 2A-2B) MitoSox intensity was similarly increased indicating increased ROS in the mitochondria as well. (Fig 2C-2D)

**Figure 2.**
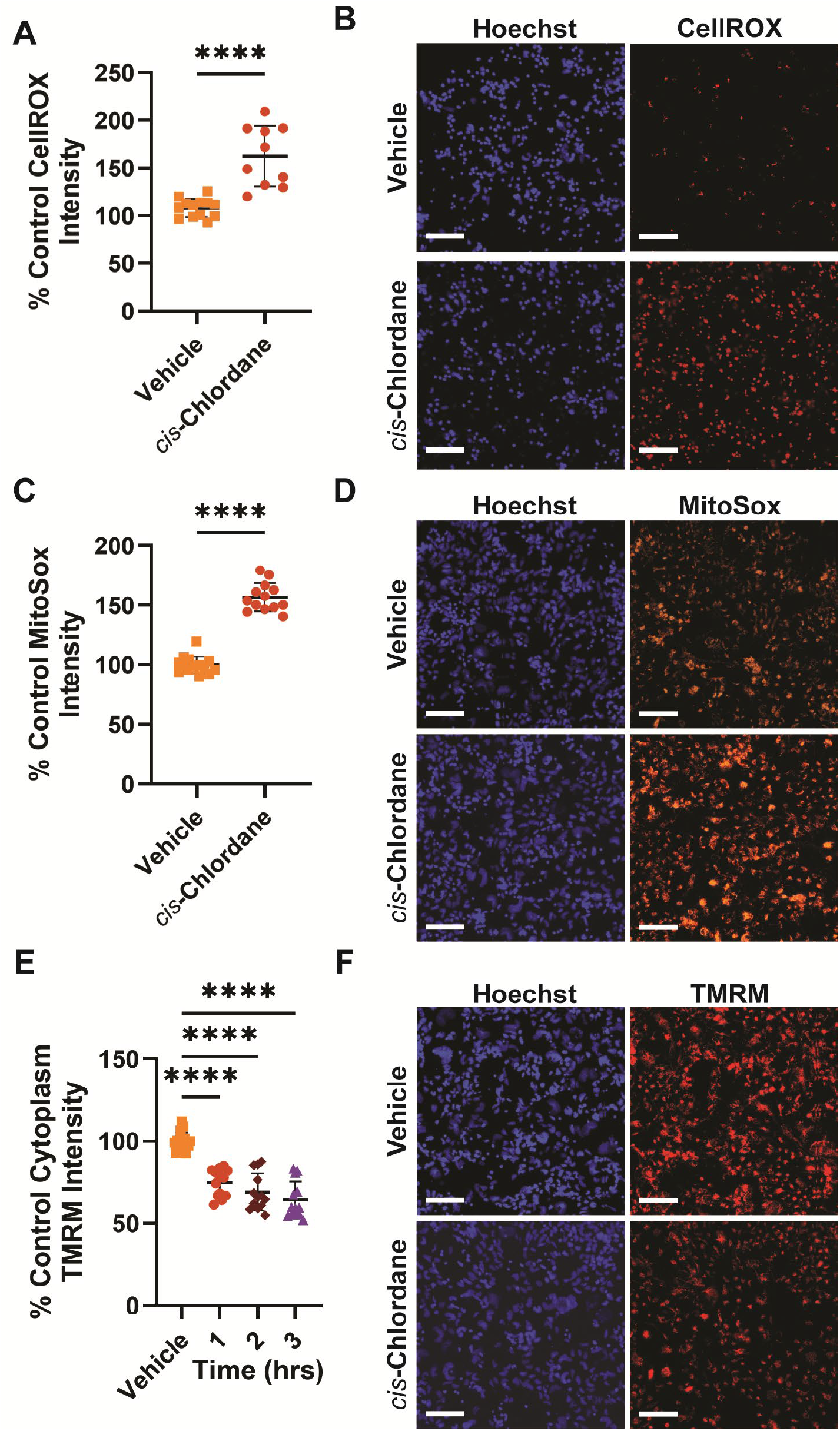
*Cis*-chlordane treatment causes increased cellular and mitochondrial reactive oxygen species and disrupts mitochondrial membrane potential in motor neurons. (A) Differential cellular ROS production in motor neurons treated with *cis*-chlordane. (B) Representative CellRox images. (C) Differential mitochondrial ROS production in motor neurons treated with *cis*-chlordane. (D) Representative MitoSox images. (E) Differential mitochondrial membrane potential in motor neurons treated with *cis*-chlordane. (F) Representative TMRM images. Bars indicate mean with SD. Statistics were performed by unpaired student T-test or one-way ANOVA with multiple comparisons to control, ****p<0.0001. Scale bar represents 100µm.

### 2.4. *cis*-Chlordane treatment disrupts mitochondrial function

To probe the integrity of mitochondrial membranes in motor neurons treated with *cis*-chlordane, we performed additional imaging using the dye TMRM, which accumulates in mitochondria with intact membrane potential. After only 1 hour, motor neurons treated with *cis*-chlordane bore significantly reduced TMRM intensity in comparison to control, with 2 and 3 hour treatments showing progressively more decreased signal (Fig 2E-2F). These results indicate that *cis*-chlordane incurs a significant and incremental loss of motor neuron mitochondrial membrane potential.

To assess whether oxidative stress in motor neurons treated with *cis*-chlordane correlates with altered metabolic activity, we performed a Seahorse Cell Mito Stress Test, allowing for the acquisition of multiple metabolic parameters including oxygen consumption rate (OCR), extracellular acidification rate (ECAR), and ATP consumption. Strikingly, upon 3 hours of either 10 μM or 5 μM cis-chlordane treatment, motor neurons demonstrated significantly decreased OCR, ECAR, ATP production, spare respiratory capacity, non-mitochondrial respiration, maximal respiration, and proton leak (Fig 3).

**Figure 3.**
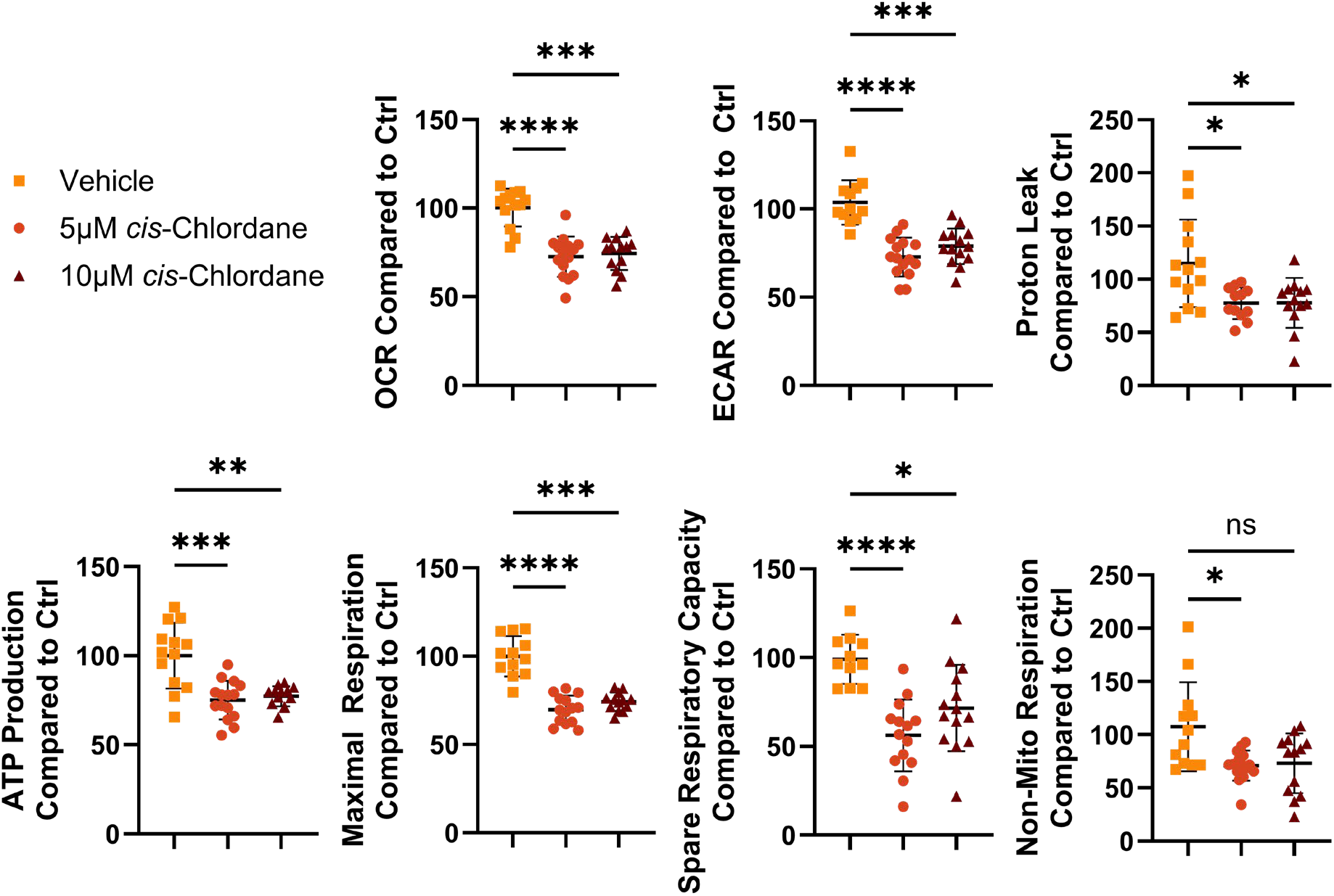
Motor neuron metabolism is highly altered upon treatment with *cis*-chlordane. Motor neurons were treated with either 5μM *cis*-chlordane, 10μM *cis*-chlordane, or vehicle (DMSO) for 3 hours before performing Agilent’s MitoTox XFe Seahorse assay. Values were normalized to total DNA present in subsequent lysates, and treatment values then compared to in-plate control. Bars indicate mean with SD. Statistics performed with Kruskal-Wallis test with multiple comparisons. *p<0.05, **p<0.01 ***p<0.001, ****p<0.0001

Interestingly, we performed a pilot of the Seahorse Cell Mito Stress Test using HEK293T cells treated with the same amount cis-chlordane and for the same amount of time, and found the opposite effect on metabolism. In HEK293T cells, basal OCR, ATP production, maximal respiration, spare respiratory capacity, proton leak, and non-mitochondrial respiration, are all significantly *increased*. (Fig S5) This suggests that the mechanisms or extent of toxicity mediating HEK293T and motor neuron susceptibility to *cis*-chlordane is different.

To further probe for metabolic changes, we performed luminescence-based assays measuring pyruvate and malate concentrations in *cis*-chlordane treated motor neurons. While pyruvate (product of glycolysis) content remained unchanged from controls, malate (a metabolite in the citrate cycle and part of the malate-aspartate shuttle) content in cell lysates was significantly increased following 3-hour treatment with *cis*-chlordane (Fig 4). In comparison, when the same assessments were performed on *cis*-chlordane-treated HEK293T cells, neither pyruvate nor malate concentration was altered in comparison to vehicle control. (Fig S6)

**Figure 4.**
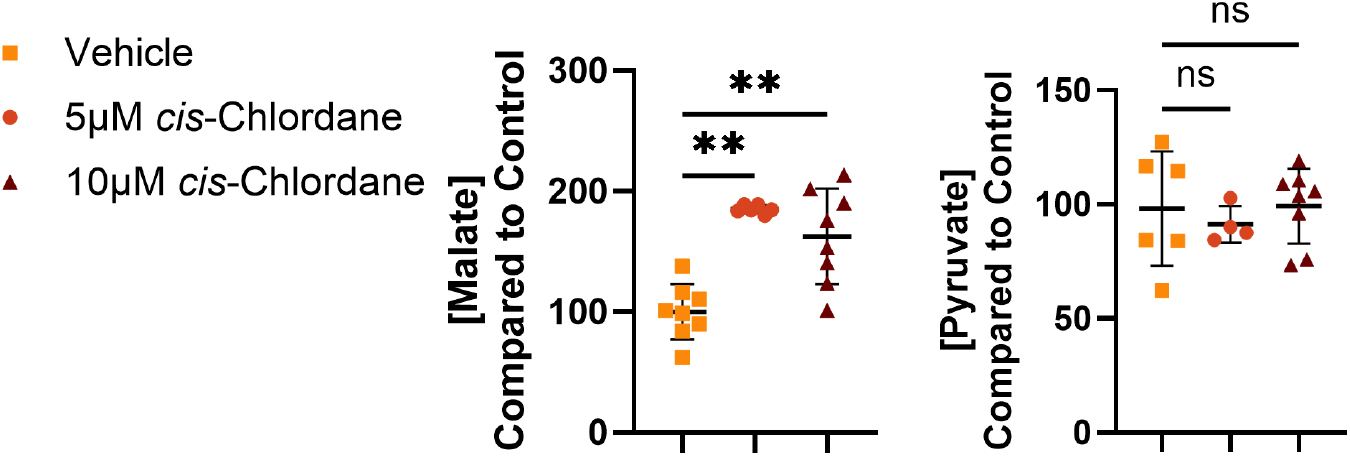
Concentration of metabolic intermediates in *cis*-chlordane treated motor neuron is altered. Motor neurons were treated with *cis*-chlordane or vehicle for 3 hours before the cells were lysed and quantitatively assessed for Pyruvate or Malate. Measured values were compared to in plate control. Statistics performed through one-way ANOVA with multiple comparisons to control. ***p<0.001, ****p<0.001

## 3. Discussion

Herein, we establish novel modes of toxicity associated with the organochlorine pesticide *cis*-chlordane, demonstrating significant ALS-like mitochondrial perturbations in exposed motor neurons. Postmortem assessments have revealed oxidative damage in ALS patient motor cortices [24], and we demonstrate a highly altered oxidative stress profile in motor neurons treated with *cis*-chlordane, including increases in both mitochondrial and overall cellular ROS. It is well documented that ROS production in the mitochondria is highly dependent on membrane potential [25]; indeed, we also observe a significant and progressive loss of membrane potential in chlordane-treated motor neurons, suggesting electron transport chain (ETC) uncoupling. Interestingly, this ETC uncoupling is not due to increased proton leak; in fact, motor neurons treated with cis-chlordane experience reduced proton leak. Loss of membrane potential in chlordane-treated motor neuron mitochondria may thus be attributed to dysfunction in one or multiple ETC complex proteins, causing reduced proton efflux into the intermembrane space and subsequent limitation of leakage. Supporting this notion, we observed significant decreases in both OCR and ATP production following *cis*-chlordane treatment, indicative of decreased complex IV and ATP synthase activity. As such, specifying the effects of *cis*-chlordane on each ETC complex will be an immediate topic of study. Defects in ETC complexes I, II, III, and IV have been observed in postmortem ALS patient cortical tissue [26], and downstream oxidative stress is highly implicated in ALS pathogenesis. The data presented here suggest that *cis*-chlordane treatment induces a similar mitochondrial phenotype in stem cell-derived motor neurons, adding further complexity to the pesticide’s toxilogical profile. The opposite metabolic response of HEK293T and motor neurons to *cis-*chlordane suggests not only a unique mode of toxicity in either cell type, but highlights the importance of testing chemicals in the correct cell type that matches the disease of interest.

In addition, we observed significant buildup of malate, but not pyruvate, following *cis*-chlordane treatment, possibly contributing to downstream ETC malfunction. The mitochondrial inner membrane is impenetrable to NADH, resulting from the citrate cycle and the main electron carrier of the ETC, and therefore relies on the malate-aspartate shuttle to pass the reducing equivalents into the mitochondrial matrix. While altered malate content has not been characterized in ALS patients, in SOD1 models of ALS, the malate-aspartate shuttle is implicated as a mechanism of metabolic pathogenesis, influencing both citrate cycle and oxidative phosphorylation processes [27]. Similar dysfunction may connect the phenotypes observed here, compounding cellular malate concentration and impairing electron transport (in the form of reducing equivalents/NADH) into the ETC.

Mitochondrial hypopolarization and subsequent ROS generation have been observed in SOD1 models of ALS [28, 29], where mutant SOD1 aggregation characterizes cellular phenotypes. However, when we employed a protein aggregation assay, no changes in protein aggregate content were observed between cis-chlordane-treated motor neurons and control. These findings contrast our RNA-sequencing results, which suggested increased expression of ER stress genes and downstream UPR proteins upon 16 hours of pesticide exposure. ER stress has emerged as a candidate for the pathogenesis of sporadic ALS [30], with one canonical ALS hallmark being high quantities of ubiquitinated inclusions throughout the nervous system, often bound to the autophagosome-localizing P62 [31].

Autophagy, another mode of cellular ER stress amelioration, was significantly upregulated as indicated through RNA-seq, and an increase in acidic endosome content was observed through fluorescence microscopy after *cis-*chlordane treatment. While this may indicate that ER stress initiated by *cis*-chlordane is primarily ameliorated by autophagic mechanisms, the observed vesicle buildup may also be the product of increased mitophagy and improper mitochondrial flux. While aberrancies in mitochondrial turnover have been demonstrated in mutant models of ALS [32], the specificities underscoring neuronal mitophagy regulation are still unknown. As such, investigating whether these vesicles are indeed mitophagic will be a topic of future study.

In conclusion, our findings further implicate *cis*-chlordane as an inductive factor in the development of ALS phenotypes in motor neurons. In particular, *cis*-chlordane produces a metabolic phenotype in motor neurons bearing similar dysfunction to both canonical ALS models and ALS patient tissue samples. We propose that the *cis*-chlordane-induced mitochondrial phenotype increases ROS accumulation in the cell, in turn damaging proteins and initiating the observed ER stress/UPR response. Understanding the primary origin of pesticide-induced mitochondrial perturbations will be an immediate focus in future experiments.

## 4. Materials and Methods

### 4.1. Pesticides

Cis-chlordane (Agilent, Santa Clara, USA) was resuspended in DMSO (Amresco, Solon, USA). Multiple lots of *cis*-chlordane were resuspended with very similar levels of toxicity; EC50’s ranged from 12-16µM.

### 4.2. Cell Culture

Differentiation into motor neurons was performed as previously established [33], using the WT 1016a iPSC line (Harvard Stem Cell Core, Cambridge, USA) or the WT hESC line WA09-Islet-GFP [34]. Following differentiation, motor neurons were aliquoted at 20*10^6^ cells/mL in a liquid nitrogen tank. Cells were thawed in complete media containing Neurobasal, 1X N2, 1X B27, 1X penicillin streptomycin, 1X NEAA, 1X Glutamax, 3.2 mg/mL glucose, 0.2 mM ascorbic acid, 10 ng/mL BDNF/GDNF/CTNF (Gibco, Eugene, Oregon) and plated in black 96-well plates (Greiner, Monroe, USA) at a density of 30,000 cells per well. Plates had been coated with poly-D-lysine (Corning, Glendale, USA) and poly-orthine (Fisher Scientific, Eugene, USA) in 100mM borate buffer pH 8.3 overnight, then laminin (2.5 µg/mL) and fibronectin (5 µg/mL) (Invitrogen, Eugene, USA) overnight. Cells were incubated for 72hr after thawing before media were replaced with complete media containing pesticide or vehicle only. Representative images of the WA09 and 1016a motor neurons with and without *cis-*chlordane treatment can be found in Fig S7.

HEK293T cells were plated in Dulbecco’s Modified Eagle Medium (DMEM) (Invitrogen, Eugene, USA) containing 10% Fetal Bovine Serum (Gibco, Eugene, Oregon), 1X Penicillin-Streptomycin (Gibco, Eugene, Oregon), 1X GlutaMAX (Gibco, Eugene, Oregon), and 1X Non-Essential Amino Acid Supplement (Gibco, Eugene, Oregon). For experimental procedures, HEK293T cells were plated in black 96-well plates at a density of 5,000 cells/well. Cells were incubated for 24 hours after plating before the media was replaced with complete DMEM containing pesticide or vehicle only.

### 4.3. Fluorescence Microscopy

After an accordant incubation period in media containing *cis*-chlordane, cells were stained with 10μg/mL Hoescht (Invitrogen, Eugene, USA) and either 10μM CellROX Deep Red or 1μM MitoSox in oxidative stress assessments (Invitrogen, Eugene, USA), 75nM LysoTracker Deep Red in lysosome content assessments (Invitrogen, Eugene, USA), 250nM TMRM during membrane integrity assessments (ThermoFisher, Eugene, USA), or 50μg/mL Propidium Iodide during viability assessments (Invitrogen, Eugene, USA). After incubation at 37°C for 30 minutes, plates were imaged live using a ImageXpress Pico Automated Cell Imaging System (Molecular Devices, San Jose, USA) using automated capture at 10x magnification. Subsequent image analysis was performed in CRX.

For Lysotracker staining, average positive cell fluorescence intensity was normalized to in-plate control wells and statistical significance determined using a Kruskal-Wallis test with multiple comparisons. For CellROX and MitoSox staining, average positive cell fluorescence intensity was normalized to in-plate control wells and statistical significance determined using an unpaired T-test. For TMRM staining, average positive cell cytoplasmic fluorescence intensity was normalized to in-plate control wells and significance determined using an Ordinary one-way ANOVA with multiple comparisons to control.

### 4.4. RNA sequencing

WT WA09-Islet-GFP human stem cells were differentiated into motor neurons as described previously and frozen in a liquid nitrogen storage tank. This cell line expresses GFP when the transcription factor Islet1, a marker of motor neurons, is expressed. Cells were thawed and allowed to grow for 72 hours before media was changed to complete media containing 16μM cis-chlordane or vehicle only (DMSO). After 24 hours of treatments, cells were FACS sorted on a BD FACS Aria II Cell Sorter (BD Biosciences, Franklin Lakes, NJ), and RNA isolated using TRIzol (ThermoFisher, Eugene, USA) according to the standard protocol. RNA quality was assessed with an Agilent Tapestation (Agilent, Santa Clara, USA) and sent for mRNA sequencing to the University of Connecticut Center for Genome Innovation on an Illumina NovaSeq6000 (Illumina, Coyoacán, Mexico) producing single-end reverse strand 100bp reads.

Raw sequencing reads were downloaded from Illumina Basespace as fastqsanger.gz files and uploaded into Galaxy to perform an RNA-seq analysis pipeline [35]. The sample libraries for replicates run in separate lanes were concatenated into individual technical replicates using Concatenate datasets [36]. Quality of the raw sequencing reads was checked using FASTQC [37] with no unexpected biases or failures found. Overrepresented sequences were checked using NCBI BLAST [38], with no matches found in the human genome, indicating that no contaminants were present in the samples. Reads were mapped to the built-in hg38 human genome on Galaxy using HISAT2 [39] with default settings for single-end reverse strand reads, and the genome (GTF) file available at the time on ensemble-Homo_sapiens.GRCh38.103.gtf.gz [40]. Processing, statistical analysis, and visualization was done using R (in Rstudio) (R Development Core Team, 2010; Rstudio Team, 2019) using iDEP (v 0.93) resources and techniques [41].

Fold changes, false discovery rates (FDRs), and DEGs were identified using DESeq2, with an FDR cutoff of p < 0.05 and a minimum fold change of 2 used to identify DEGs. GO term enrichment analysis was performed on DEGs using the Gene Ontology Consortium online database [42]. Similarly, generally applicable gene-set enrichment (GAGE) with the KEGG database was done using the GAGE R package [43] with the KEGG online database [44]. Genes with an FDR > 0.5 were removed prior to analysis, and the enrichment FDR cutoff for significance was set at p < 0.05. Absolute values of fold changes were used for disease pathway enrichment analysis. Significant patways of interest were visualized on KEGG pathway diagrams using the pathview R package [43].

### 4.5. RT-qPCR Validation of RNA-seq

After 24 hours of treatment with 16 µM of *cis*-chlordane, GFP-Islet1 motor neuron cultures were sorted with FACS and the RNA isolated using TRIzol (ThermoFisher, cat # 15596026) according to the standard protocol, with an extra isopropanol precipitation and ethanol wash. cDNA of isolated mRNA was synthesized using a OneScript® Plus cDNA Synthesis Kit (Applied Biological Materials, cat # G236) with Oligo(dT)s. qPCR was performed with a total volume of 20 µL/well with the following volumes/concentrations: 20 ng of cDNA, 10 µL of Blastaq 2x qPCR master mix (Applied Biological Materials, cat # G891), and 0.5 µM F/R primers from Eurofins Genomics (see primer sequences in Table S1) on a BioRad CFX 96 instrument. The recommended Blastaq qPCR Master Mix fast protocol was used with two notable changes: the activation step increased to 3 min, and the annealing step/extension temperature decreased to 57°C, as multiple primers had melting temperatures of approximately 62°C.

### 4.6. Proteasome Activity Assay

After 24hr treatment with 10 μM cis-chlordane, 1016a motor neurons were washed with PBS and lysed with a 10-minute incubation with 1X RIPA lysis buffer, Benzonase, and 1mM PMSF. Lysates were centrifuged and a Pierce Rapid Gold BCA Protein Assay (ThermoFisher, Eugene, USA) performed to measure lysate protein concentration. 20 μg of lysates were transferred to a black 96-well plate and 100 μL of proteasome activity buffer (Abcam, Waltham, USA). MG132 (Selleckchem, Houston, USA), a known proteasome inhibitor, was included as a positive control. Fluorescent substrate was then added to a final concentration of 50 μM. Samples were incubated for 1 hour before assessed for fluorescence (Ex/Em 350/440 nm) using a M5 plate reader (Molecular Devices, San Jose, USA). Statistical testing was performed in Prism using an Ordinary one-way ANOVA with multiple comparisons to control.

### 4.7. PROTEOSTAT Protein Aggregation Assay

After 3 hours of treatment with either 10 μM *cis*-chlordane, 5 μM *cis*-chlordane, or vehicle control, cells were lysed in RIPA buffer. Lysates were transferred to a 384-well plate and manufacturer instructions were followed for the PROTEOSTAT Protein Aggregation Assay (Enzo, Farmingdale, NY), utilizing 40% of the recommended volume for reagents to account for the miniaturized assay format. During assay protocol, wells were assessed for fluorescence (Ex/Em 550/600 nm) using a Varioskan Lux Multimode Microplate Reader (Thermo Scientific, Waltham, USA). Significance was determined in Prism using an Ordinary one-way ANOVA with multiple comparisons to control.

### 4.8. SeaHorse XF Cell Mito Stress Test

Motor neurons were plated at 200,000 cells per well and HEK293T cells were plated at 50,000 cells per well in an Agilent XFe 24 well plate and allowed to adhere for 72 hours. They were then treated with 5 and 10 µM *cis*-chlordane in fresh media for 3 hours at 37°C. The plate was then subjected to Agilent’s MitoTox XFe Seahorse assay following the manufacturer’s instructions. After the run, the wells were lysed and total DNA content was used to normalize each well to the amount of cells present. Three biological replicates with 4 technical replicates each were run. Data was compared to in plate control averages. The in plate control-normalized data was then compiled across biological replicates and significance determined through either Kruskal-Wallis test with multiple comparisons or unpaired student T-test.

### 4.9. Promega Pyruvate-Glo and Malate-Glo Kits

1016A iPSC-MNs were plated and allowed to grow for 72 hours prior to experimentation. Following 3hr incubation in fresh media containing 10 μM or 20 μM *cis*-chlordane, cells were lysed using the reagents included in either the Promega Pyruvate-Glo, or Malate-Glo kits (Promega, Madison, USA) in accordance with manufacturer recommendation. Following completion of the included protocol, lysates were transferred to white 96-well plates and assessed for luminescence using a Varioskan Lux Multimode Microplate Reader (Thermo Scientific, Waltham, USA). Significance was determined using Dunnett’s multiple comparisons tests in Prism, or unpaired Student T-test for comparisons involving just two treatment groups.

## Supplementary Materials

Supplementary Fig files include: Volcano plot of DEGs, GO and KEGG analysis, ProteoStat data, HEK293T SeaHorse and metabolic data, representative motor neuron images, and qPCR primer sequences.

## Author Contributions

Conceptualization of study, A.O.; methodology, A.O.; formal analysis, O.C.; investigation, O.C., M.H., A.W.; writing—original draft preparation, O.C.; writing—review and editing, O.C., A.O.; visualization, O.C.; supervision, A.O.; funding acquisition, A.O. All authors have read and agreed to the published version of the manuscript.

## Data Availability Statement

The full RNAseq dataset has been uploaded to GEO, accession number TBD.

## Acknowledgments

We would like to thank Wesleyan University’s Bailey College of the Environment for their continued support. We would also like to thank Wesleyan University Professor Joseph Coolon for his assistance in analyzing our RNAseq data.

