## Supplemental Information for "Exposure to the organochlorine pesticide cis-chlordane induces ALS-like mitochondrial perturbations in stem cell-derived motor neurons"


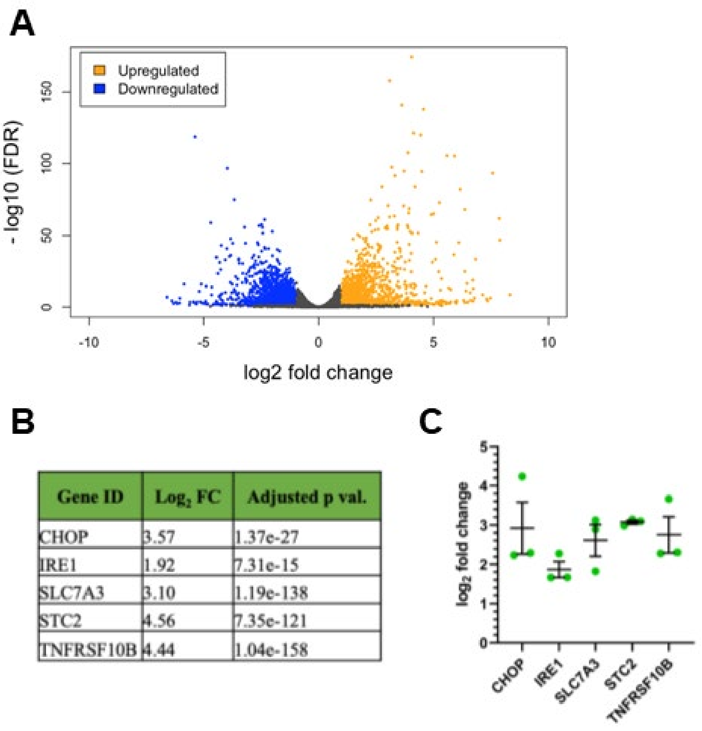


| Gene ID | $Log_2$ FC | Adjusted p val. |
|---|---|---|
| CHOP | 3.57 | 1.37e-27 |
| IRE1 | 1.92 | 7.31e-15 |
| SLC7A3 | 3.10 | 1.19e-138 |
| STC2 | 4.56 | 7.35e-121 |
| TNFRSF10B | 4.44 | 1.04e-158 |

**Fig S1. DEGs in motor neurons treated with *cis*-chlordane. A)** Analysis with DESeq2 indicated 1764 genes were significantly upregulated (orange) and 1806 genes were significantly downregulated (blue) in response to 16 hours of *cis*-chlordane treatment. Increased values on the y-axis indicate higher significance. The cutoff values for colored DEGs were the product of the FDR value cutoff on y-axis and the fold change cutoff of 2 on the x-axis. **B)** Table of representative highly upregulated genes from RNAseq data. **C)** qRT-PCR of the genes in B confirming sequencing results in a freshly prepared sample of hESC motor neurons treated with cis-chlordane.

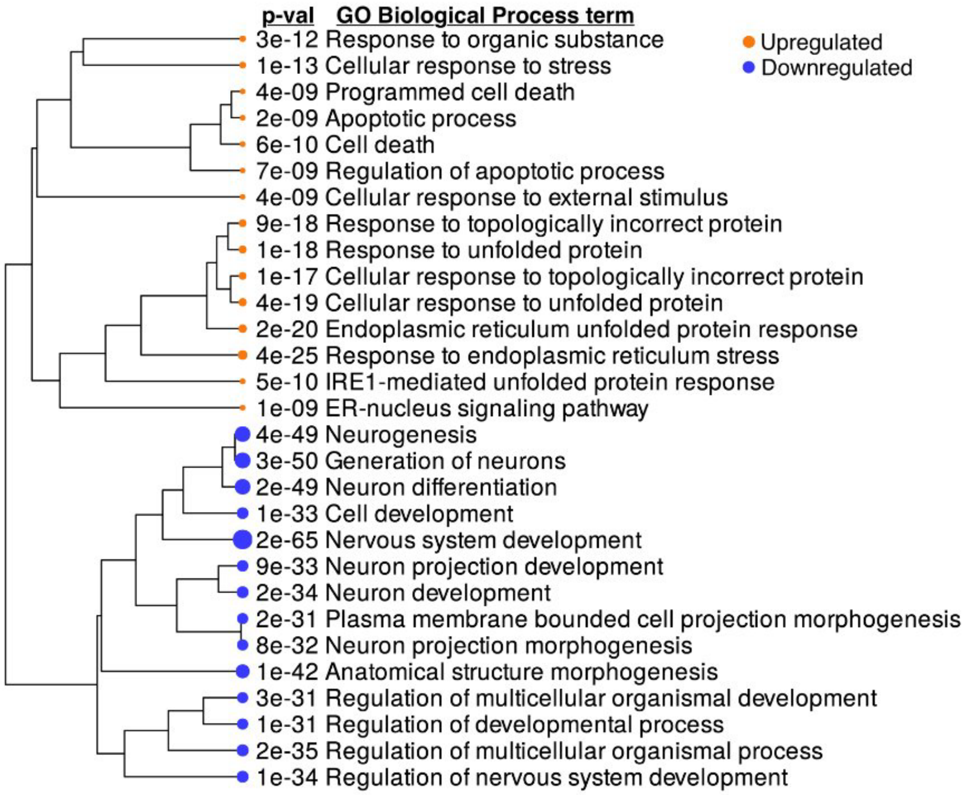


**Fig S2. Gene Ontology (GO) analysis of differentially expressed genes as determined by RNA-sequencing of motor neurons treated with *cis*-chlordane.** Generally applicable gene set enrichment (GAGE) analysis was performed on RNA-seq data, allowing for the determination of pathways with altered expression. The top 15 most significant enriched terms for upregulated and downregulated groups of genes are shown. Dot size is directly related to significance, where a larger dot indicates a more significant p-value, shown to its right.

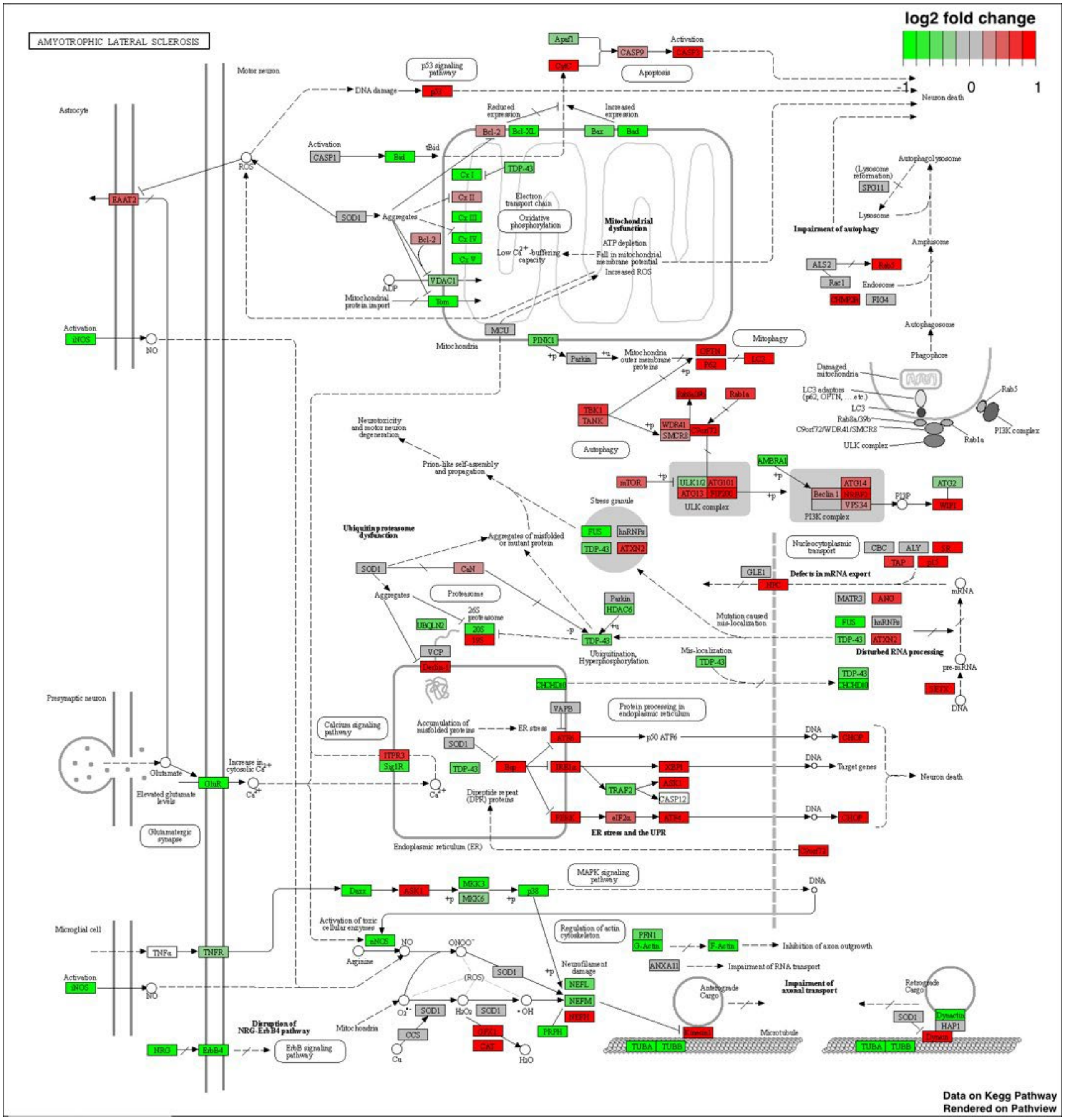

**Figure S3. Motor neurons treated with *cis*-chlordane show transcriptomic changes similar to those in ALS.** RNA-seq data was aligned with the “ALS” KEGG disease pathway, indicating differential expression of ALS-implicated genes.

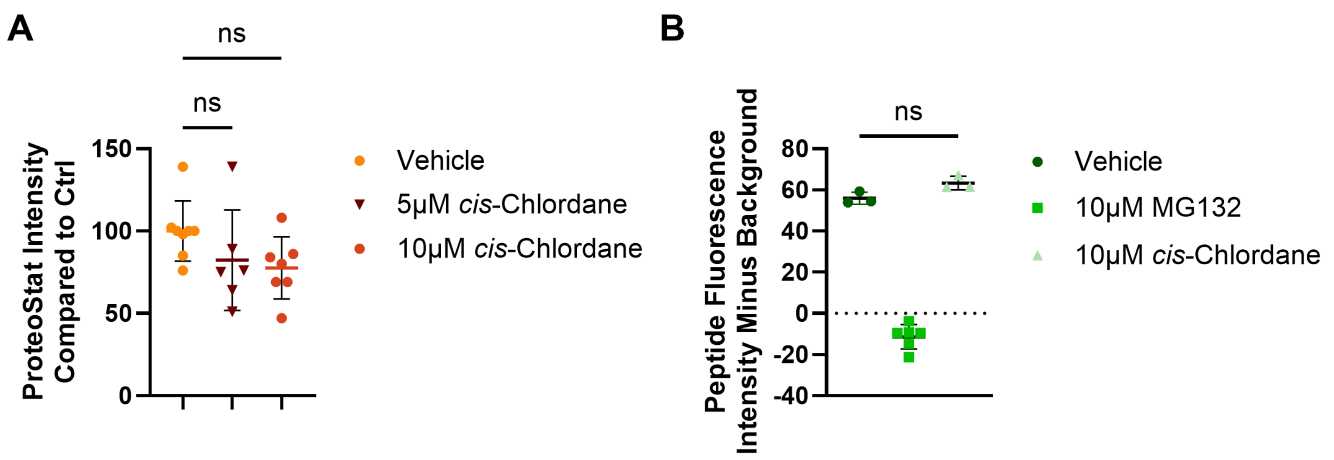


**Figure S4. Motor neurons treated with *cis*-chlordane demonstrate no changes in protein homeostasis compared to control. A)** PROTEOSTAT output detailing changes in protein aggregate content between treatment groups. Motor neurons were treated with *cis*-chlordane or vehicle control (DMSO) for 3 hours before execution of assay protocol. Treatment fluorescence was compared to in-plate control average. **B)** Changes in proteasome activity associated with *cis*-chlordane treatment. Motor neurons were treated with *cis*-chlordane, vehicle control (DMSO), or the proteasome inhibitor MG132 for 24 hours before being lysed. Fluorescent substrate was then added to lysates and fluorescence quantified.

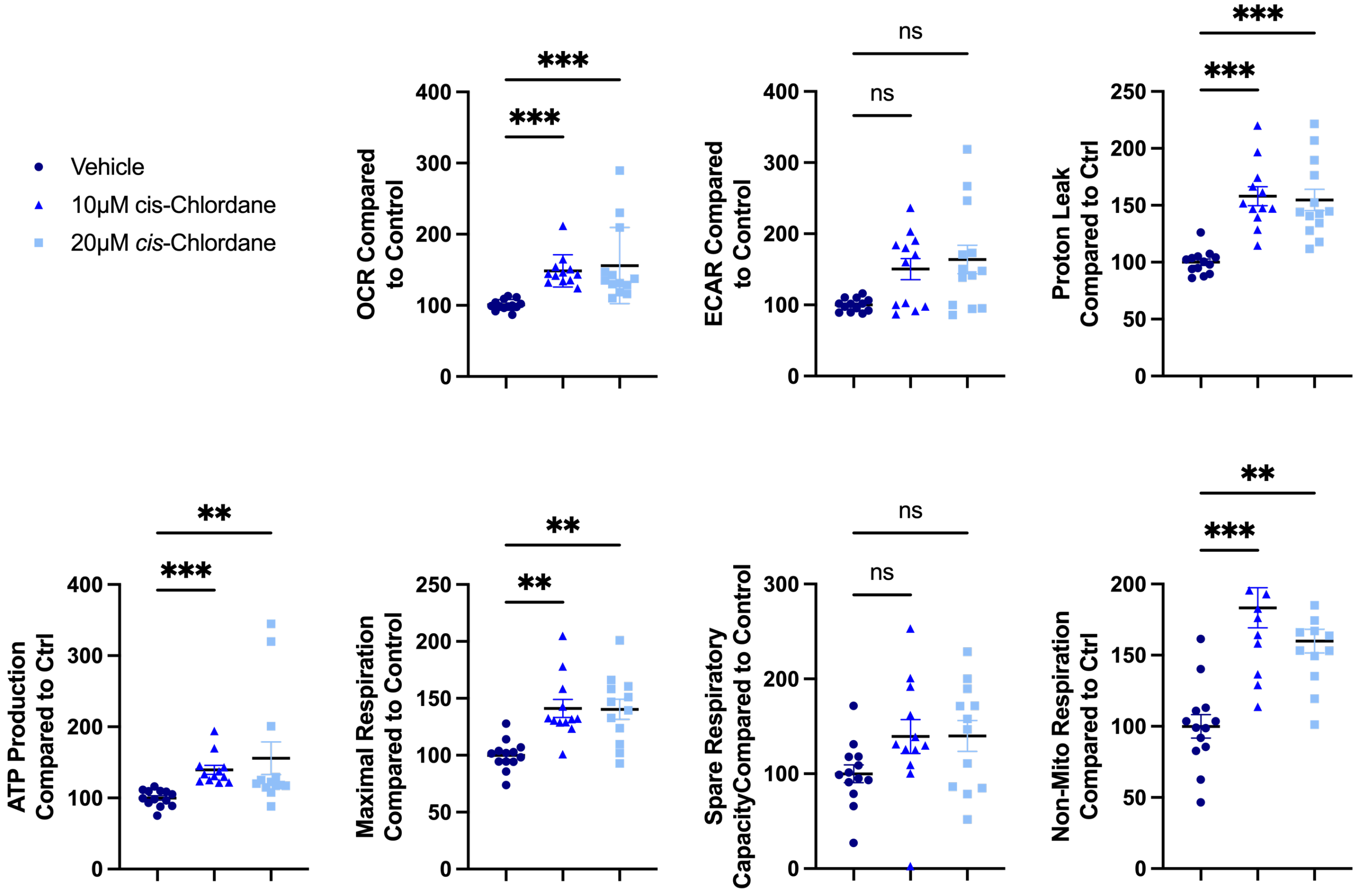


**Figure S5. HEK293T metabolism is altered in response to *cis*-chlordane treatment.** HEK293T cells were treated with either 5μM *cis*-chlordane, 10μM *cis*-chlordane, or vehicle (DMSO) for 3 hours before performing Agilent's MitoTox XFe Seahorse assay. Values were normalized to total DNA present in subsequent lysates, and treatment values then compared to in-plate control. Bars indicate mean with SD. Statistics performed with Kruskal-Wallis test with multiple comparisons. *p<0.05, **p<0.01 ***p<0.001, ****p<0.0001

### HEK293T

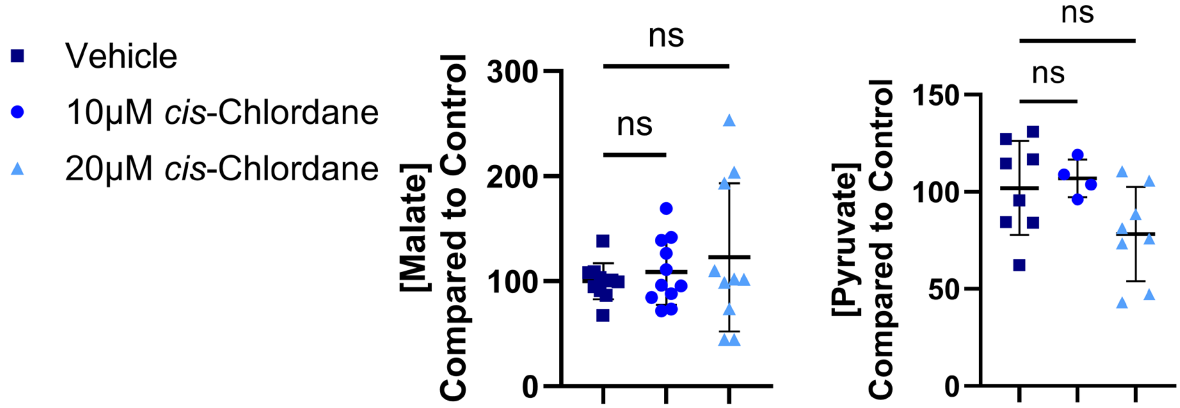


**Figure S6. Concentration of metabolic products in *cis*-chlordane-treated HEK293T cell lysates.** HEK293T cells were treated with *cis*-chlordane or vehicle control for 3 hours before cells were lysed in accordance with Pyruvate-Glo or Malate-Glo protocol. Following protocol execution, luminescence was acquired and RLU values compared to in-plate control. Statistics performed through one-way ANOVA with multiple comparisons to control. ***p<0.001, ****p<0.0001.

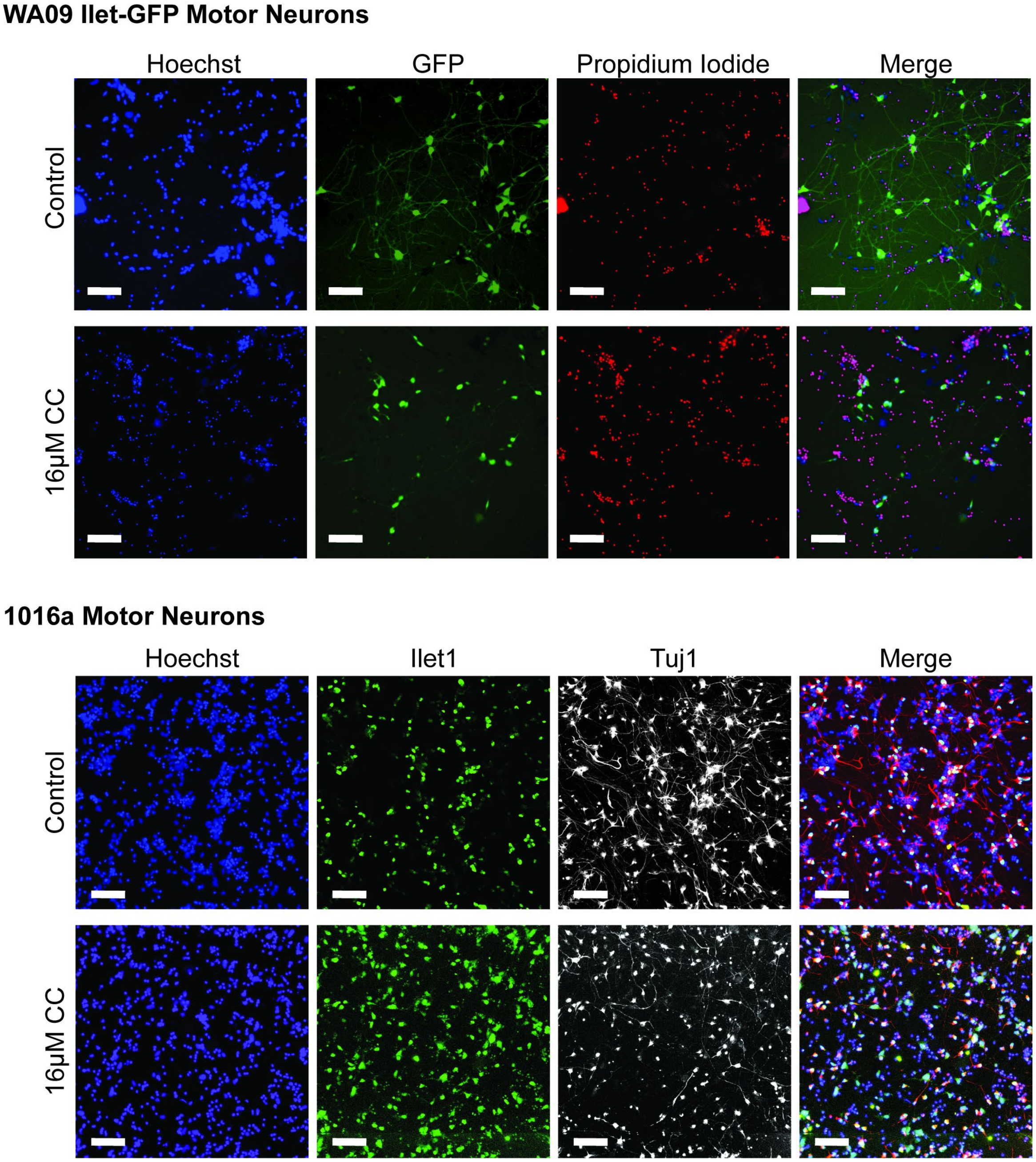


**Figure S7. Representative images of human stem cell derived neurons treated with cis-chlordane.** WA09-Islet-GFP is a human embryonic stem cell line that expresses GFP under the motor neuron specific Islet1 promoter resulting in "free floating" GFP that fills the neuron. The WA09 motor neurons were imaged live at 10x with Hoechst (nuclei) and propidium iodide (PI, dead cells) counter stains. In the *cis*-chlordane treated WA09 neurons, notice the increase in PI stain and marked decrease in neurite complexity. 1016a is a wild-type human induced pluripotent stem cell line. After treatment, the cells were fixed in 4% PFA and immunostained for motor neuron nuclei (Islet1), neurites (Tuj1, beta-tubulin), and nuclei (Hoechst) as described in the Methods and imaged at 10x. In the *cis*-chlordane treated 1016a neurons, notice the marked decrease in neurite complexity. Both samples were treated for 24 hours with 16μM of *cis*-chlordane. Scale bar represents 100μm.

**Supplemental Table 1:** qPCR primer sequences used to verify RNAseq data.

GAPDH F: 5’-AAGGTGAAGGTCGGAGTCAAC-3'

GAPDH R: 5’-GGCGTCATTGATGGCAACAATA-3'

CHOP F: 5’-GATGAAAATGGGGGTACCTATG-3'

CHOP R: 5’-AGGGCTAACATTCTTACCTCTTCA-3'

IRE1 F: 5’-GAAGCATGTGCTCAAACACC-3'

IRE1 R: 5’-TCTGTCGCTCACGTCCTG -3'

SLC7A3 forward, 5′-CACTCAACTCCATCCCCACT-3′

SLC7A3 reverse, 5′-CTGTGGCTGTCTCCAGATGA-3′

STC2 forward, 5′-ATGCTACCTCAAGCACGACC-3′

STC2 reverse, 5′-TCTGCTCACACTGAACC TGC-3′

TNFRSF10B forward: 5′-GCCCCACAACAAAAGAGGTC-3′

TNFRSF10B reverse: 5′-AGGTCATTCCAGTGAGTGCTA-3′
